# INDIVIDUAL CONTRIBUTIONS TO NEST BUILDING AND INCUBATION BY COOPERATIVELY BREEDING AMERICAN CROWS

**DOI:** 10.1101/2024.02.01.578444

**Authors:** Carolee Caffrey, Charles C. Peterson, Tiffany W. Hackler

## Abstract

American Crows (*Corvus brachyrhynchos brachyrhynchos*) are long-lived birds with pair bonds that may last many years. Pairs in Stillwater, OK, nested singly or in groups with up to 10 auxiliaries (Caffrey and Peterson 2015). Breeders did most of the nest building. Pair members contributed at approximately equal rates, although the sexes differed slightly in details: males tended to carry sticks and hand off materials more often, and females spent more time at and in nests. Both sexes worked faster on second and third attempts than on first attempts of the season. We found no evidence that pair members were (sexually) signaling to each other via their contributions. Incubation periods were characterized by low levels of activity at nests, where females spent most of their time and were fed once every 3-4 hours, mostly by their mates. Contributions to both stages of nesting by auxiliaries varied widely and exhibited no patterns with respect to any measured phenotypic characteristics.

## INTRODUCTION

Nest building is costly behavior (Moreno et al. 2010, Mainwaring and Hartley 2013), and several authors have posited sexually-selected signaling benefits as drivers of nest-builders’ investment decisions (e.g., Soler et al. 1998, Gill and Stutchbury 2005, Moreno 2012, Mainwaring and Hartley 2013). Most such thinking, however, is couched in scenarios of breeding with new mates; to our knowledge, few ideas have been published regarding what might be expected of the effort committed to this stage of annual reproductive attempts by long-term mates in long-lived birds. If year-to-year survivorship and the likelihood of breeding together again are both high, some form of long-term budgeting by breeders of both sexes is likely to underlie decisions regarding current investments.

American Crows are long lived, and many breed over successive years with the same mates. In such relationships, pair members might be expected to “work things out” to the best advantage of both, which presumably would mean fledging and thereafter successfully raising an optimum number of healthy offspring each year. As such, we were interested in examining details of the contributions of females and males to the building of nests, to shed some light on this behavior in a bird with long-term, socially-monogamous pair bonds. From our experience watching crows build nests and incubate eggs over many years, we predicted females and males would contribute relatively equally to building, with females spending more time attending to nest interior details.

We examined, too, the behavior of auxiliaries (Caffrey and Peterson 2015) in the contexts of nest building and incubation. Given the tendency for individual crows to vary widely in behavior independent of commonly-measured phenotypic characteristics (Caffrey and Peterson 2015, Caffrey et al. 2016b), we did not expect to find any significant patterns to auxiliary contributions to these two stages of cooperative breeding attempts.

## METHODS

### Background

Our study population of marked crows in Stillwater, Oklahoma, had been under observation since August 1997, and is described in Caffrey and Peterson (2015). Individual crows were captured as free-flyers (Caffrey 2002a) or obtained as nestlings, and all individuals were marked with patagial tags and unique combinations of colored plastic and U.S. Fish and Wildlife Service metal leg bands (Caffrey 2002b and c). Crows caught as free-flyers were aged via inspection of plumage and mouth color pattern characteristics (Emlen 1936); adults were assigned the age “≥2 y” at marking. For the present study, crows were observed during the nest-building and incubation stages of the breeding seasons of 2001 and 2002. Observations were made with binoculars and spotting scopes, primarily from vehicles, except for those of one group (#29 in 2001; Table 1), whose placement of their nest next to a window in an empty building allowed occasional continuous videotaping of nest activities for up to many hours at a time. (A blind next to the window provided our only means of observing this extremely wary group. Unfortunately, we were repeatedly locked out of the building, and so were unable to observe Group 29 on many days.)

**Table 1.**
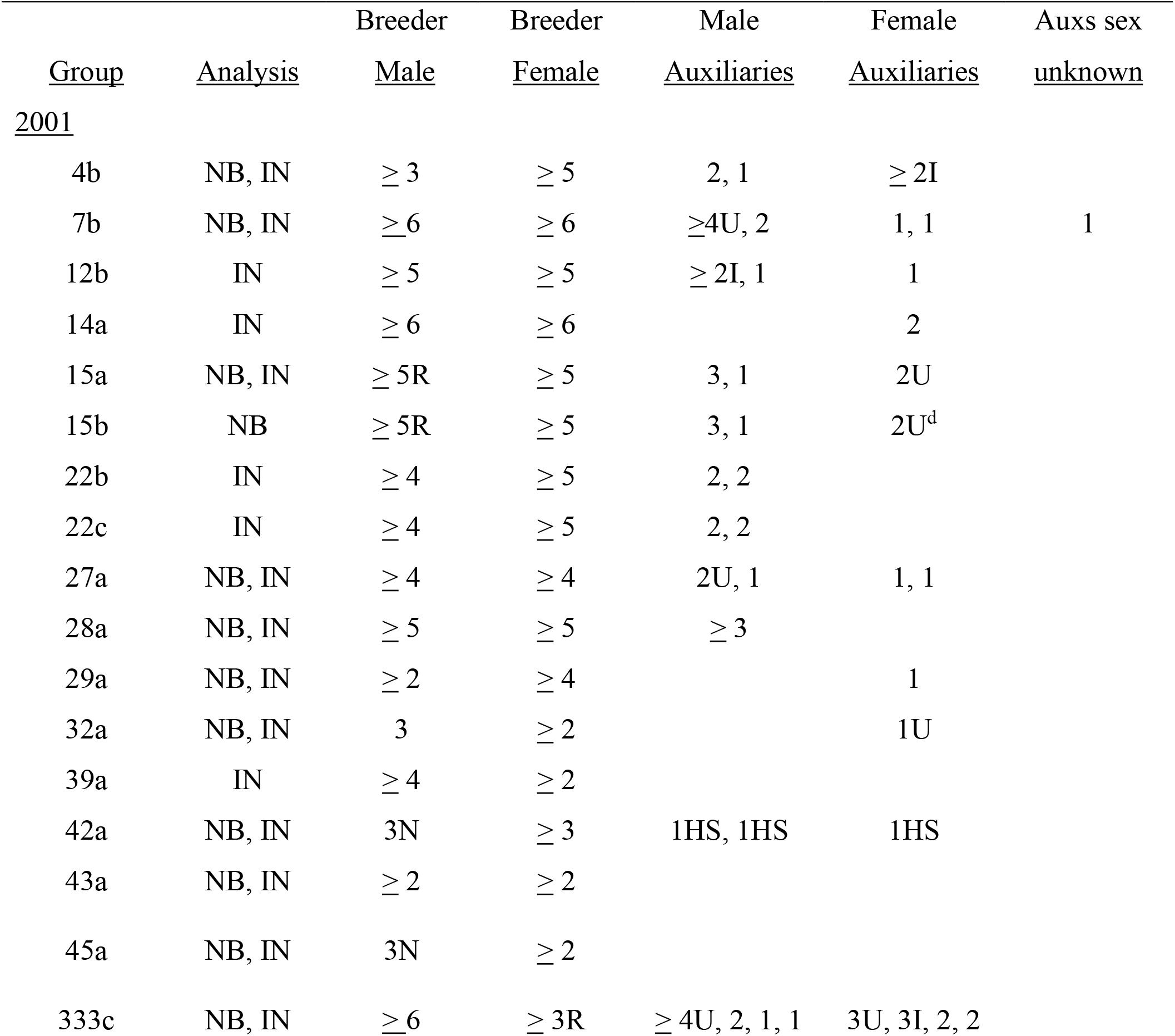
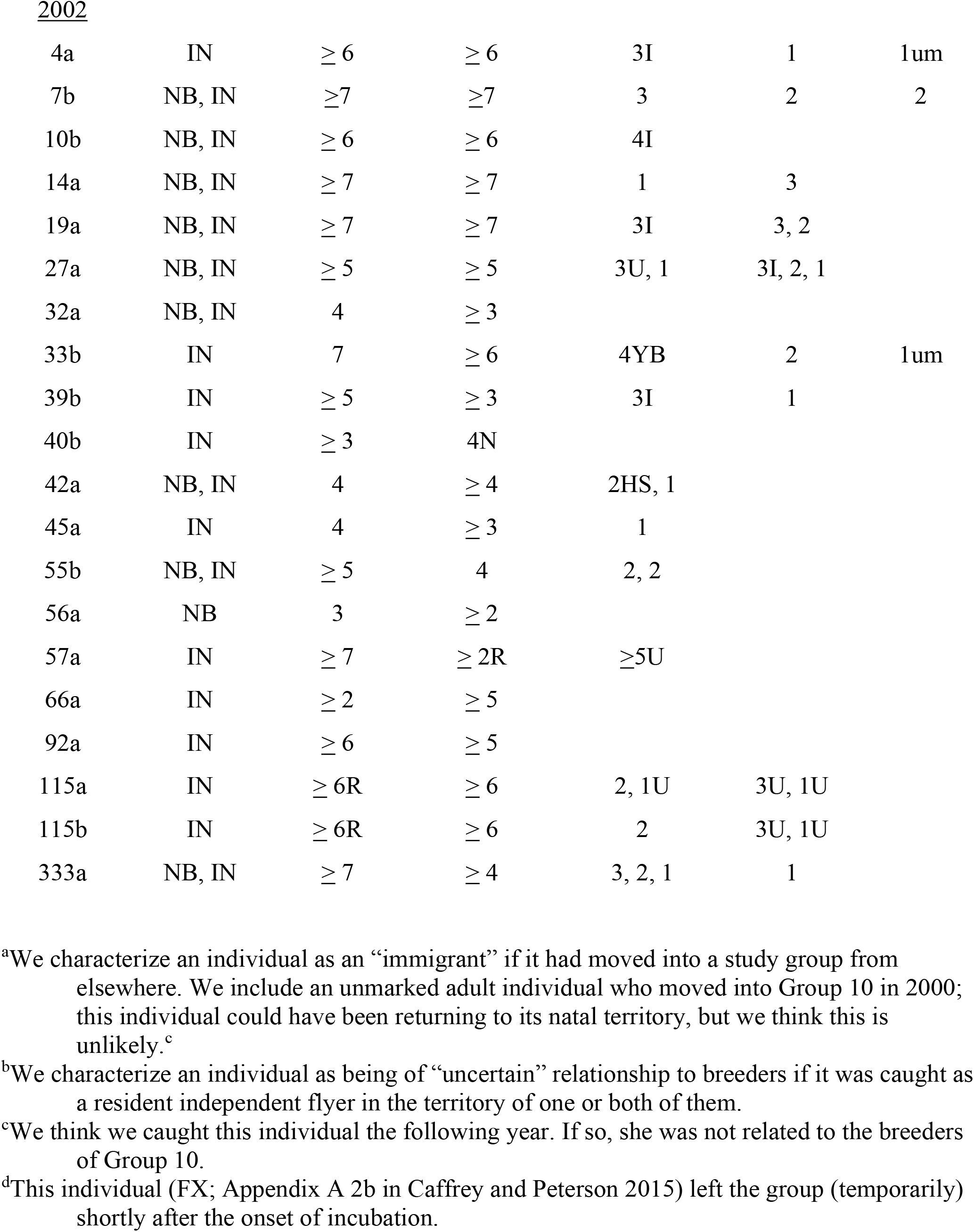
Composition of crow groups included in nest building (NB) and incubation (IN) analyses. Each numeral represents an individual by its age in years. Most auxiliaries had hatched in nests of one or both breeders. N (novice): known to be breeding for first time. R: breeder was replacement of social parent of at least one of auxiliaries in group. I: immigrant^a^. HS: half-sib of male breeder. YB: younger brother of male breeder. U: dispersal history (relative to group) uncertain^b^. um: unmarked.

### Genetic Sampling and Analyses

We collected approximately 1cc of blood from the brachial vein of each crow for molecular determination of sex and for estimation of microsatellite-based relatedness values, analyzed parentage in the program Cervus (Kalinowski et al. 2007), and estimated microsatellite-based relatedness coefficients (r) in the program Relatedness (Queller and Goodnight 1989). Procedural details can be found in Caffrey and Peterson (2015).

We did not have DNA samples for one adult female auxiliary (in Group 333 in 2001; “3U”) and four auxiliaries whose sex was unknown (Table 1). Throughout this paper, where appropriate, we report r values as means ± 95% CIs.

### Individual Contributions to Nesting Attempts

We made detailed observations of the nesting activities of individual crows. No group contained more than one unmarked individual. Nest visibility was enhanced by clipping branches at three nests in 2001 and five in 2002.

### Nest Building

Nest-building watches were distributed throughout the day from late February through mid-April. The majority of nests were located by following crows carrying nesting material. We report nest-building data from watches of nests that were completed and ultimately went on to incubation: 12 nests by 11 groups in 2001 (Table 1; 27 watches; mean = 74.7 min) and 10 by 10 in 2002 (Table 1; 35 watches; mean = 93.7 min). We identify first, second, and third nesting attempts (within years) as “a,” “b,” and “c.” During nest-building watches, we recorded trips and the materials (sticks or lining) carried by individuals, whether materials were worked into nests or handed off to other crows, whether individuals got into nests, and the total time spent at nests by each individual.

### Incubation

We determined the timing of the start of incubation primarily by observing the behavior of female breeders; see Results. We closely monitored groups as their nests approached completion, checking nests repeatedly throughout appropriate days for evidence that females had begun incubating eggs. Our monitoring included repeated brief checks in nest areas, observation periods that lasted 20-30 minutes, and nest-building watches, as described above. We also checked many nests at dusk during these days, and females settled in nests at dark were considered to have started incubation on that day.

Incubation watches were distributed throughout the day from mid-March through early May; we conducted 66 at 16 nests (15 groups) in 2001 (mean = 95.3 min [including four videotaped watches at Nest 29 for 220, 350, 240, and 480 min]), and 132 at 19 nests (18 groups) in 2002 (mean = 94.3 min). During incubation watches, we recorded trips to nests by individuals, documented their activities (e.g., whether or not visitors fed or attempted to copulate with incubating females), and determined the amount of time individuals spent at and in nests.

### Analyses

Data from nest watches were statistically analyzed for patterns related to individual- and nest-level variables. Rates of nest building changed over the nest-building period (Fig. 1a and b), and therefore a potential source of bias was the inevitably different timing of watches across nests. Because some individuals were observed to work on nests for only subsets of nest-building periods, direct comparison of building rates would be confounded by such unequal temporal sampling. In an attempt to remove this potential source of bias, we constructed overall quadratic regressions of nest-building rates on day of nest-building period and its square, and then calculated residual rates from the equation for each individual during each watch. Individuals were then represented by their average residuals over their entire sampling period for statistical analyses.

**FIG. 1.**
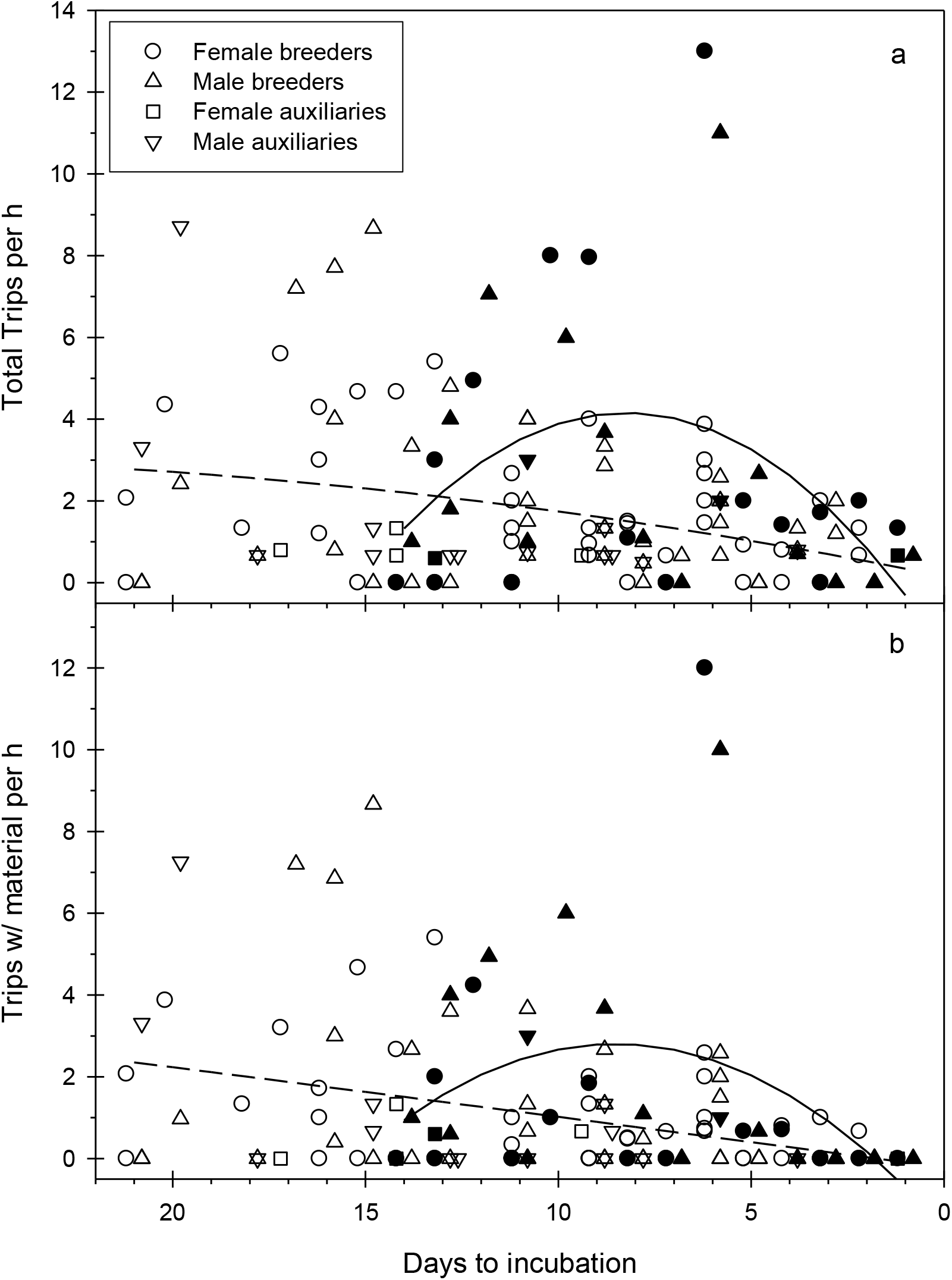
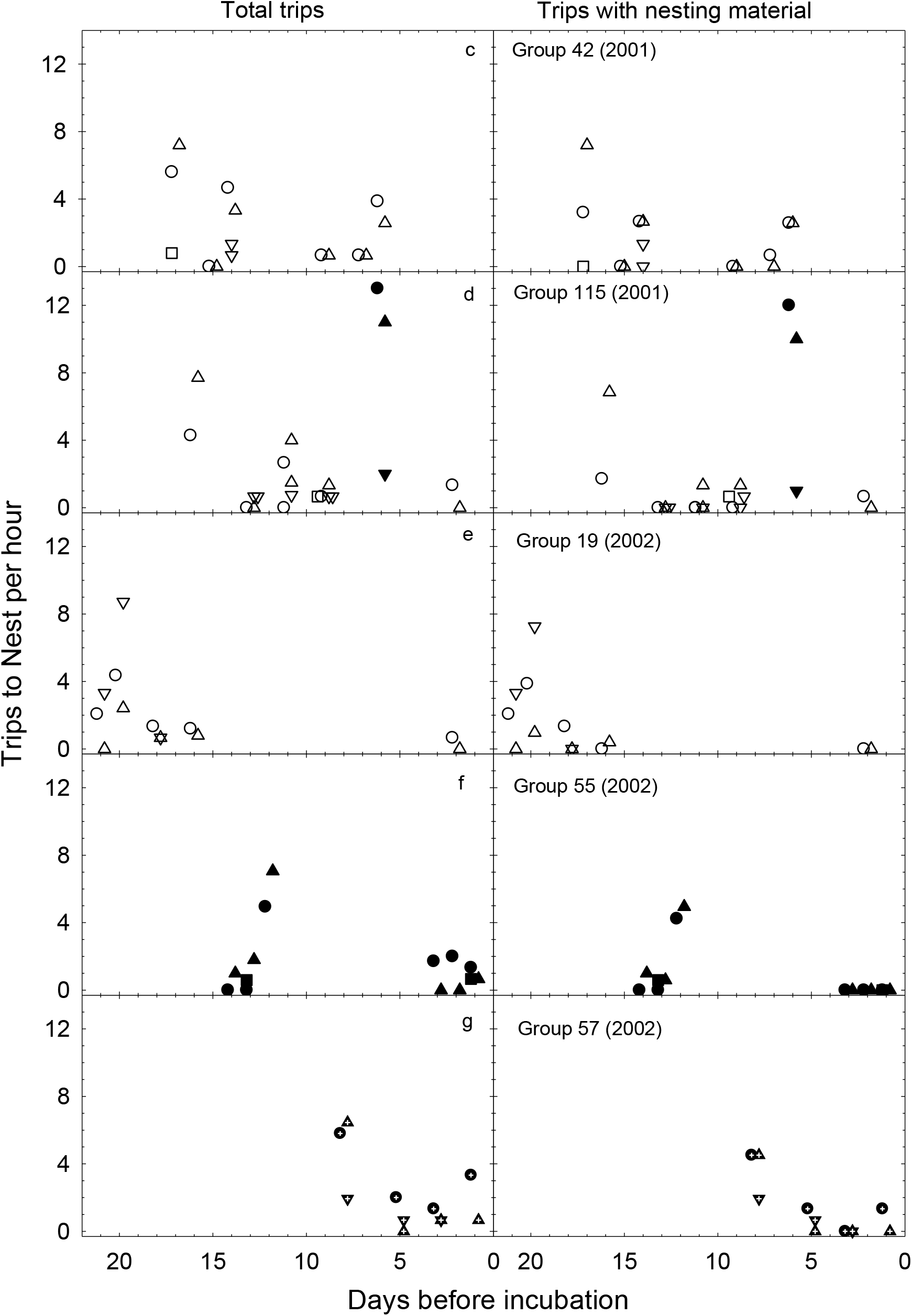
Trips to nests per hr by breeding and auxiliary crows during nest building in 2001 and 2002. Unfilled symbols are first-attempt data; filled are those of second and third attempts. Some data are displaced laterally for clarity. (a) Total trips to nests/hr. Dashed curve is best-fit quadratic for first attempts; solid is that for subsequent attempts. (b) Trips/hr in which sticks or lining material carried to nests. (c) A common pattern: crows began nests 14-17 days prior to incubation and took 7-10 days to build them. Shown are data from the first (successful) attempt of Group 42 (Appendix A 1a in Caffrey and Peterson 2015) in 2001. Visits by male breeder’s three younger half-siblings: sister didn’t contribute material but got in and arranged sticks, one brother briefly inspected from rim, one brother made two trips with material, got in and worked into nest. During the five days prior to the start of incubation, group members sometimes seen but rarely in nest area. (d) First and second attempt data; second attempt late in season (second week of April, after an abandoned first attempt wherein incubation had begun; Appendix I 2b). Visits to nest by nonbreeders 14 and 11 days prior to onset of incubation made by males from a neighboring group (Appendix A 2d in Caffrey and Peterson 2015). Two separate watches done on Day 11. Three nonbreeders (two males) at nest on day 9 were auxiliaries. Second nest completed in about four days. (e) The pair (#19) and adult immigrant male auxiliary (GN; Appendix A 2g in Caffrey and Peterson 2015) were found working on a nest 21 days prior to the start of incubation; within a week they were done. During the following week, group members (=5) were sometimes observed foraging and loafing in various areas of their territory, including near the nest tree, but sometimes could not be found, and were never observed at the nest. During the subsequent week, the male breeder and GN were each seen to briefly visit the nest once; the female breeder made infrequent brief visits, sometimes getting in. During the last pre-incubation watch for this group (=90 min in late afternoon), the female made only one visit to the rim and the male was not observed at the nest. (f) Second attempt but still early in the season (mid-March). Nest completed in about six days. During subsequent week, pair and two male auxiliaries (Group 55; Appendix A 1b in Caffrey and Peterson 2015) sometimes near, at, and in nest, and sometimes gone from territory. (g) Group 57 in 2002 (Appendix A 2c in Caffrey and Peterson 2015): third attempt of male breeder and likely first of his replacement mate, first week of April. This unclassifiable group was not included in nest-building analyses.

We treated nesting attempts in different years as independent. A variety of preliminary analyses showed no differences between years and so data from the two years of this study were combined for statistical analyses of behavioral rates and time spent at nests.

Correlation, regression, and ANOVA analyses were conducted with use of Microsoft Excel and the General Linear Model module of SYSTAT software. Our field data were characterized by wide variation (Fig. 1), and our purposes in testing statistical hypotheses are largely heuristic. We therefore apply a liberal criterion of statistical significance, discussing all results with a probability of Type I error (P) ≤ 0.1. Means, including those of regression residuals, are shown ± 95% Confidence Intervals.

## RESULTS

### Group Composition

In 2001 and 2002, group size ranged from 2-10, and group composition was highly variable (Table 1). Intra- and inter-group social dynamics were complex, the result of the variable behavior of individuals; examples are provided in Appendix A 1 and 2 in Caffrey and Peterson (2015).

Three individuals were known to be breeding for their first time in 2001 or 2002 (Table 1, Appendix I 1).

### Nest Building

Nest-building activities were observed from March 1 through April 9 in 2001 and February 22 through April 6 in 2002. In 2001, 11 study groups completed 12 nests (in which eggs were incubated): eight first attempts, three second attempts, and one third; one group (#115) renested following the abandonment of their first attempt (after incubation had begun), and is included in both “a” and “b” data sets. In 2002, seven first attempts and three second attempts by 10 groups went on to incubation. (The building of an eleventh nest, the third attempt of a male whose “replacement” mate’s recent history was unknown [Group 57 in Appendix A 2c in Caffrey and Peterson 2015] is discussed in Appendix I 2a, below, and is depicted in Fig. 1g, but was not included in nest-building analyses [Fig. 1g Caption]).

Crows nested in a variety of tree types, especially conifers and deciduous hardwoods, and usually placed nests 16-22m high in central locations in solid branch crotches (Appendix II; data from the year 2000). On several occasions we observed crows “trying out” different locations with first sticks and then moving on to other sites. Once nest bases were in place, crows continued adding sticks and began working on lining interiors as soon as nest cups began to take shape. Completed crow nests in Stillwater had exterior diameters of about 50 cm and were about 23 cm tall (n=6 nests, CC unpubl. data), and were lined with a variety of materials, including (primarily) bark shredded by breeders, other soft vegetation, mud, and mammal fur. We have also found string, ribbon, corrugated and other packing material, fabric-softening dryer sheets, a man’s tube sock, and a pair of men’s white briefs(!) incorporated into nest linings.

Most pairs began work on first-attempt nests (n=15) approximately 14 to 17 days prior to the onset of incubation and took about 7 to 10 days to complete them (Fig. 1a and b), although nest-building patterns were variable (Fig. 1c-e). Second- and third-attempt nests (n= 7) were completed more quickly (Fig. 1a, b, d, f, g).

Of 426 total trips to nests observed in 88.3 hours of watches beginning 21 days prior to onsets of incubation, 418 could be attributed to particular individuals. Breeders made 362 of 418 trips (87%; Fig. 1a), and worked on nests for 0.25-5 minutes during visits whether building material was carried or not. Breeders arrived alone or together (they sometimes waited for each other nearby), and carried nesting material (sticks or lining) on 224 trips (Fig. 1b); on 40 of these occasions the material was transferred to a mate already present. Some crows, after handing off carried material, worked elsewhere on the nest. Breeders got into nests on 248 of 362 trips.

We analyzed time-adjusted nest-building efforts of breeders with two-way ANOVA. Overall and per hour, female and male breeders did not differ in the total number of trips they made to nests (main effect of sex: F_1,40_=0.037, P=0.85; sex x attempt interaction: F_1,40_=0.026, P=0.87) or the number of trips made with building material (sex: F_1,40_=0.057, P=0.81; sex x attempt: F_1,40_=0.006, P=0.94). Breeders of both sexes tended to work faster on second or third attempts (N=7) compared to first attempts (N=15; main effect of nest attempt: total trips/h: F_1,40_=3.62, P=0.064; trips with material/h: F_1,40_=2.257, P=0.14), and breeders of both sexes, especially females, spent more time at second and third attempt nests during building than those of first attempts (nest attempt: F_1,40_=3.852, P=0.056, attempt x sex interaction: F_1,40_=2.499, P=0.121). Female and male breeders differed slightly in their nest-building contributions: of approximately equal numbers of trips to nests (188 and 174 for females and males, respectively), females and males carried lining material on 28 and 29% of them but males carried more sticks (40% of trips *vs*. 27% for females). Females and males worked on nests during 62 and 56% of trips, respectively, males tended to hand off the material they carried more often than females (17% *vs*. 6%, of trips), and females tended to get into nests more often than males (78% *vs*. 58% of visits). Overall, females spent more time at and in nests than males during nest-building periods (Fig. 2; means = 6.5 *vs*. 3.6 min/h; F_1,40_=5.283, P=0.026).

**FIG. 2.**
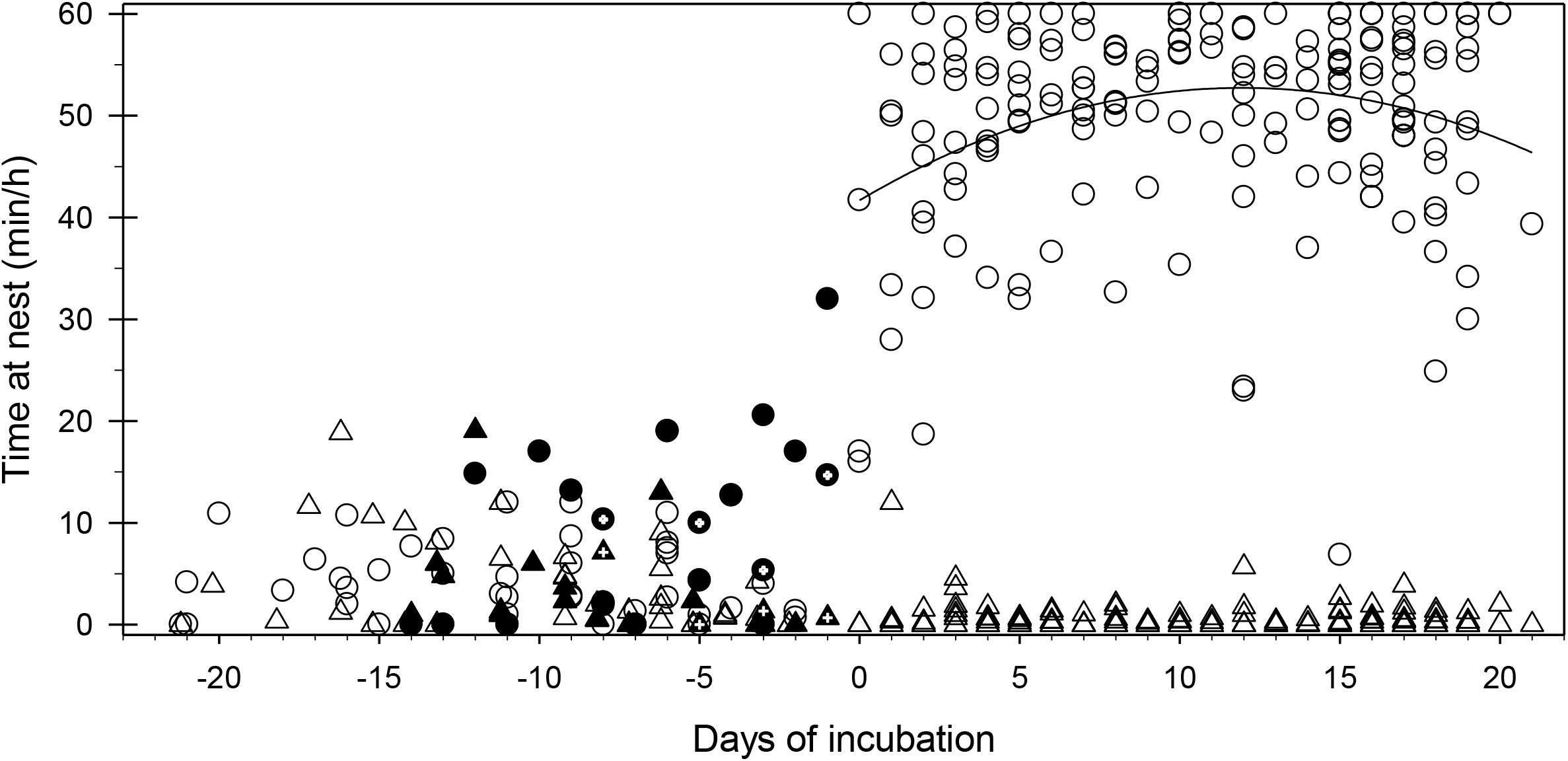
Time spent at nests during nest building and incubation by breeding crows in 2001 and 2002; sex and status symbols as in Fig. 1. Prior to incubation (to the left of zero on horizontal axis), unfilled symbols are first nesting-attempt data; filled are those of second and third attempts (cross-hatched are those of Group 57 in 2002; Fig. 1g, Appendix I 2a, and Appendix 2c in Caffrey and Peterson 2015). Data for all attempts (first and subsequent) pooled to right of zero on X axis (during incubation). Curve is best-fit quadratic.

The 56 total trips to 11 nests by nonbreeders during nest-building watches were made by 17 individuals (including two individuals in both years). A single 2-year old immigrant male auxiliary (GN; Appendix A 2g in Caffrey and Peterson 2015) made 29 of the 56 trips by nonbreeders, carrying nesting material on 23 of them (days 21, 20, and 18 pre-incubation, Fig. 1a, b, and e). Other nonbreeders that went to nests during watches included 14 other auxiliaries (yearling and adult females and males) and a yearling and adult auxiliary (both males) from a neighboring group that on one occasion together visited a nest while group members were not present; the adult (TM) returned two days later while the male breeder was nearby (Appendix A 2d in Caffrey and Peterson 2015). Of the 25 trips to nests by auxiliaries other than GN, material was carried on 18 (and often worked into nests); these were made by yearling and adult female and male auxiliaries, one of which was an immigrant (TM). The other seven trips, made by a yearling female and two yearling and four adult male auxiliaries, involved inspections from rims and within nests. Extra-pair yearlings and adults of both sexes also got into and sat in nests, including the adult male from a neighboring group (TM).

As first-attempt nests were completed, many breeders and auxiliaries disappeared from nest areas (and territories) for periods ranging from several hours to a day or two at a time, for 2-5 days in a row. As incubation approached, group members were more regularly seen and breeders spent time near, at, and in nests (Fig. 2, and Discussion: Nest Building). Within a day or two of the onset of incubation, and during the first days thereof, some females produced quiet “whine” or “waah” vocalizations (Verbeek and Caffrey 2002) while in nests.

In 2001 and 2002, five marked auxiliaries (one in both years) built nests separate from those of the breeding pairs in their groups: Appendix I 3.

### Incubation

We report data for 15 groups and 16 incubation attempts in 2001, and 18 groups and 19 attempts in 2002; two groups (#22 in 2001, #115 in 2002; Table 1) renested after failed first attempts (after hatching had occurred) and we include both sets of watches in our data set.

We defined the start of incubation as the day female breeders met one or more of the following criteria: they were settled in nests during several brief checks throughout the day, they remained motionless in nests for extended periods during nest watches or one or more 20-30 minute checks/day, or they were settled in nests at dark. (The presence of male breeders in guarding positions [Discussion: Incubation] was used as corroborative evidence.) In two cases we recorded incubation as beginning despite the low percent of time female breeders spent in nests during watches (Fig. 2, Day 0) because in both cases females left nests early in watches and did not return until later in the time allotted, at which point they settled low into nests. Both females were in nests when checked later those same days. Females began incubating first attempts as early as March 10 and as late as April 6 in 2001, and March 8 and March 24 in 2002. Incubation dates for second- and third-attempts that later hatched eggs were as late as April 22 in 2001 and May 2 in 2002.

Only female breeders incubated eggs. During incubation, they spent the majority of time in nests (Fig. 2), leaving infrequently mostly to stretch, defecate, or forage. In multiple regression, the amount of time females spent in nests depended on the day of the incubation period (*t*_172_ = 3.90; P< 0.001) and its square (*t*_172_= 3.50; P = 0.001), as well as on time of day (*t*_172_= 2.00; P = 0.047) and its square (*t*_172_= 2.42; P = 0.016), such that females were most likely to be in nests during the core days of incubation periods (Fig. 2), and earlier than later in the day throughout. Time spent in nests during incubation by females was not related to incubation date, nesting attempt (a, b, or c), length of incubation period, group size, or air temperature during watches.

After approximately 14-17 days of incubation, females began (and then continued) to intermittently shuffle around in nests, and to get up and put their heads down inside and do *something* that caused their tails (their only visible body parts) to shake. (Three climbs to nests revealed that eggs were moving during this period; Discussion: Incubation.) Two to four days after “first shuffle” (at most nests), the first nestling feeding trips were observed. We defined the first day of nestling feeding as the first day of hatching. This was confirmed by climbing to four nests (in days prior to and including days of first observed feeding trips) to make direct observations. Estimated incubation periods ranged from 17-21 days in 2001 and from 16-22 days in 2002, with a mean of 19 days in each year.

While incubating, females were fed at an average rate of 0.32 times per hour, with 58% of all feedings being made by male breeders. There was no difference in incubation-feeding effort between males with (N=24) and without (N=6) auxiliaries (*t*_28_= 1.56, P=0.13).

Nonbreeders made a total of 92 trips to nests in 308 hours of incubation watches. Of the 92 trips, 84 were made by auxiliaries; 43 of these (made by 25 individuals) involved the feeding of incubating females. Auxiliary patterns of incubation feeding were variable (e.g., 4 of 9 trips to a nest, 4 of 6, 2 of 4, 1 of 2, and 1 of 1 in 850, 943, 943, 872, and 767 minutes of observation, respectively) and the highest rate was low: 1 feeding trip in five hours of observation. Feeders and nonfeeders of incubating females were of varied relationships with them (Appendix 4a and b), there was no relationship between feeding rates and degree of relatedness to females in either year, and there were no differences in either year between mean relatedness values (r), to either breeder, for auxiliaries observed feeding incubating females versus those that were not (in 2001, auxiliaries that fed [n=9]: r with female breeders = 0.36±0.15, with males = 0.33± 0.11; those that didn’t [n=21]: r with female breeders = 0.38±0.09, with males = 0.27± 0.09. In 2002, auxiliaries that fed [n=14]: r with female breeders = 0.42±0.11, with males = 0.24± 0.13; those that didn’t [n=19]: r with female breeders = 0.25±0.10, with males = 0.34± 0.10).

During the 41 non-feeding trips made by auxiliaries (26 yearling and adult females and males), nest visitors would stand on rims for a few seconds up to five minutes at a time. Sometimes incubating females would get up on rims for short periods during visits and then settle back in, sometimes breeding males would join visitors on rims, sometimes females would leave for brief periods during visits, and sometimes auxiliaries would get *into* nests; sometimes while females were present and sometimes while they were not.

Two of the eight trips to nests by extra-group individuals during incubation were made by previous group members who had dispersed the previous January and February and were home for brief visits: a three-year old male spent about two minutes on the rim while the female (a replacement – two years earlier - for the original female breeder of the group) sat quietly, and a one-year old female fed her social mother and remained on the rim for two more minutes, looking around and down toward the nest, until her mother pecked at her foot and (we assume) caused her to leave. At another nest, a yearling male (ZO; a half-sib of the incubating female [AM]’s offspring with a different mate [Appendix A 1a in Caffrey and Peterson 2015), from a neighboring territory, performed a bowing display (Verbeek and Caffrey 2002) as he approached the nest rim; she snapped at him and he immediately flew away. The remaining five visits by extra-group individuals involved copulation attempts.

### Extra-pair copulation attempts

Extra-pair copulation attempts were seen on five occasions occurring from 4 to 14 days after the start of incubation. The males included two breeders, two 2-year old auxiliaries from neighboring territories and one unmarked bird of unknown identity. All five of these copulation attempts were initiated while other members of the females’ groups were not present. On one occasion, the male breeder returned during the copulation attempt and chased the intruding male out of the area; in all other cases, intruding males left on their own. Incubating females were not receptive to any of these males; they thrashed, snapped at, and clawed them, and vocalized loudly. Once intruding males departed, females returned to their positions in nests.

## DISCUSSION

In contrast to their seemingly subdued territorial behavior once nesting was underway (and for subsequent months; Appendix I 6), pairs of crows in Stillwater noisily pronounced the beginnings of breeding seasons each year by perching high and vocalizing from within territories and at peripheries. Within days, however, as pairs began working on nests, they became more secretive; so much so that, even as they flew sometimes long distances carrying nesting material (including dryer sheets, tube socks, and underwear), nest-building American Crows and their nests were (and are) difficult to find, even for seasoned searchers.

### Nest Building

Crow nests are large, intricate structures with cushy interiors that require hundreds of building trips and many hours of work to complete; for pairs building second or third nests of the season, those trips and hours were squeezed into fewer days. Female and male breeders did not differ in overall effort – making about the same average number of trips to nests per hour and carrying material in about equal proportions – but they tended to make different decisions with regard to the details of their contributions: males carried more sticks and handed off more material than did females, and females got into nests more so than did their mates. On average, females spent 80% more time at and in nests than did males (6.5 *vs*. 3.6 min/hr), and much of their time was spent fine-tuning nest interiors.

Few nest-building trips were made by individuals other than breeders, except for those made by GN (Fig. 1e, and Appendix A 2g in Caffrey and Peterson 2015), a 3-year old male who had immigrated into his group in 2001 and made 55% of nest-building trips in 2002 (Appendix 2g in Caffrey an Peterson 2015). Auxiliary trips to nests, with or without nesting material, were sometimes brief but lasted up to 14 min, and were made when breeders were present or not (Appendix I 2c and d).

Many pairs finished nests several days before females began incubation (maximum = 16 days: Group 19 in 2002; Fig. 1a and e, and Appendix A 2g in Caffrey and Peterson 2015), and in the days after nests were completed, many pairs (and group members) would be gone from nest areas and territories for long periods (several hours to 1-2 days at a time, and in the case of Group 19 in 2002, 1-2 days at a time more than once). We abandoned nest watches when groups were absent and instead spent considerable time looking for them, most times to no avail. At other times between nest completion and the onset of incubation, breeders and auxiliaries foraged and loafed in territories as if nothing was new. Group members were more regularly seen and breeders increasingly spent time near each other as incubation approached. We saw one pair copulate 4 days before the female began incubating (Appendix I 5a). Two to four days preceding the onset of incubation, females and males began spending time near and at nests; they perched side-by-side (and sometimes preened each other) or males guarded from nearby as females tended to and sat inside for one to at least 45 minutes at a time throughout the day. Females were likely laying eggs during some of these visits, as American Crows usually begin incubation with the laying of the third egg (Verbeek and Caffrey 2002) and at one nest in 2002, a climb four days pre-incubation revealed zero eggs, and two days later, one. Males guarded females closely as they foraged (either by walking and standing beside them or perching low overhead), and pair members were mostly quiet (including most of the females that produced characteristic food-begging “waahs” [Verbeek and Caffrey 2002] with increasing frequency from near and in nests) during these pre-incubation days.

Female American Crows in some populations may begin laying eggs as soon as nests are complete (references in Verbeek and Caffrey 2002) but in others females may delay egg-laying: 6-16 days in Florida and 4.8 ± 2.4 (SD) days in Saskatchewan (references in Verbeek and Caffrey 2002). The common pattern in Oklahoma of leaving nesting areas for long periods of time after ceasing work on nests was also exhibited by Western American Crows in a population in California (CC unpubl. data), and the period of days over which pairs disappeared prior to resumption of nesting activity came to be referred to as “the Poconos,” after the popular honeymoon destination of New Yorkers. We know nothing about the functions of these absences, although possibly by reducing their presence in nesting areas after a week or two of heightened activity, breeding crows might disinterest predators formerly aware of their presence (KJ McGowan, pers. comm.). Presumably breeding females were seeking nutrition in preparation for egg-laying during those times away from territories but again, we were unable to find them.

The patterns in which second and third nests were built suggest that breeders were in danger of somehow running out of time as nesting seasons progressed. Second attempts begun early in the season (mid-March) tended to thereafter follow the general pattern of first attempts; by building nests more quickly, pairs left time for trips to the Poconos (e.g., Fig. 1f). As March gave way to early and then mid-April, the honeymoons of pairs beginning new nests were cut short and incubation followed shortly after nests were completed (e.g., Fig. 1d and g, and Appendix I 2a and b). Other species of birds are known to synchronize nesting with resource availability (e.g., Blue Tits, *Cyanistes caeruleus*, and caterpillars; Helm et al. 2006), and for some species, particular resources required for normal nestling development are known (e.g., Blue Tits and taurine [in spiders]; Arnold et al. 2007). We do not know the breeding resources that might have become limiting to crows as calendar years progressed, however, the temperatures reached in typical Oklahoma summers (mean high temperature for both July and August = 34°C; usclimatedata.com) can be prohibitive to both arthropod and avian activity (e.g., Wilmer 1983 and Clark 1987, respectively).

### Incubation

For most pairs, the onset of incubation was signaled by a noticeable change in the demeanor of females – they sat deep in nests, and still, for long periods of time - and the presence of males in guarding locations in nearby high, obvious places with good views of nests and surrounding areas; males sometimes stayed in the same locations for hours at a time. Males continued guarding from above as females sat during incubation, and often stayed close to them when they left nests to stretch, defecate, or forage, especially during the first week or so, during which time we saw three within-pair copulations, all in 2002 (Appendix I 5b-d). On the second day of incubation in 2002 we also saw two *attempted* within-pair copulations (Appendix I 5e and f).

It was also during the first week of incubation (and the previous couple of days) that we saw two male breeders each actively keeping an adult male auxiliary away from nests and females: an older brother (NK) and his younger half-sib (EK, in Group 42 [2002]; Table 1, and Appendix A 1a and 2j in Caffrey and Peterson 2015), and the breeder of Group 57 and PH (his son [2002]; Appendix A 2c, and Appendix B Prediction 8, in Caffrey and Peterson 2015). Yet we saw no attempts by another eight male breeders to thwart any approaches by nine adult male auxiliaries (six immigrants, a younger brother, and two in a group with a replacement female) to either nests or females, and neither did we ever see any indication of paternity-seeking behavior by these nine auxiliaries (and yet, two of them [NX and KP] – in the same group [#333] in the two different years – both fathered young in those years; Appendix A 1b and Appendix B Predictions 8 and 12-15, in Caffrey and Peterson 2015).

The first two weeks of incubation for most groups were characterized by low levels of activity; females were mostly in nests (Fig. 2) and sat motionless for long periods while other group members mostly foraged and loafed. Females were visited at nests about once every three to four hours, mostly to be fed, and mostly by their mates. For auxiliaries that fed incubating females, the degree to which they were related to them did not influence the rate at which they fed them. Genetic relatedness to incubating females (or their mates) also did not influence the decision by auxiliaries to feed incubating females or not (during watches).

Whether visits involved feeding or not, incubating females were often completely accommodating to auxiliaries of either sex seeking peeks underneath and/or wanting to get in nests (once for almost two minutes [the whole time the female was gone] and once *with* the female). At one nest at which an (adult male) auxiliary arrived with a stick (during incubation) and began working it in, the female got up and onto the rim and waited until he finished and left before she settled back in. One female, a replacement for the social mother of a three-year old male (home for a brief visit after having moved out during the previous February [2001]), allowed him to land on the rim and hang out for about two minutes.

### Extra-pair copulations

Of the five extra-pair copulation attempts we witnessed, three were caught on videotape at Nest 29 in 2001 (Methods, and Appendix 1a in Caffrey and Peterson 2015), 13 and 14 days into incubation, in a total of 21.4 hours taped over three days (Days 13, 14, and 15 [we were unable to view this nest for the first 12 days of incubation]). The other two attempts were seen on Days 4 and 10 during totals of 12.78 and 7.18 hours of incubation watches at those nests, respectively. As above (Results), females were unreceptive to extra-pair males and fought back, in one case toppling out of the nest, flapping and holding onto the intruder with her feet; after disengaging and returning to the nest, he aggressed upon her again, and again her resistance caused the two of them to topple out. (Upon her return a second time, the male sat below her nest for about 10 seconds before flying off.)

### Concluding Remarks

Female and male members of breeding pairs collaborated on the building of nests, making approximately equal numbers of trips per hour but differing slightly in their focus: female effort was biased in favor of attending to nest interiors, presumably related to the fact that they alone incubate eggs. We found no evidence that pair members were signaling to each other via their contributions: there were no predictable relationships between nest-building and nestling-feeding rates within seasons (residual trips-with-material and residual feeding rates (Caffrey et al. 2016b): all breeders, *r*=0.32; females, *r*=0.49; males, *r*=0.29 [all P>0.10]), and there were no differences for residual time-at-nest, total trips, and trips-with-material between pairs nesting for the first time (N=5) and those (N=8) wherein mates were experienced with each other (all breeders, females, and males, P>>0.10). The lack of support for sexually-selected signaling was not surprising. Crows are long-lived and many breed with the same mates over many years. The majority of study crow pairs in our population in 2001 and 2002 had lived at least the previous year together and had just recently survived the winter, together, when nest building got underway. As the current year’s fitness interests of both sexes of breeders was presumably the same – to fledge an optimum number of healthy offspring - then, presumably, too, were their short-term interests: to efficiently build a structure capable of providing adequate warmth and protection for needy eggs and nestlings, and then to get on with subsequent stages of nesting.

As to the wide variation in, and lack of any demonstrable patterns to, contributions to nest building and incubation by auxiliary crows, this finding is itself becoming a pattern (Caffrey and Peterson 2015, Caffrey et al. 2016b).

## ACKNOWLEDGMENTS

This work was enabled by the contributions of many, and we sincerely thank, again, the people and organizations we acknowledge in Caffrey and Peterson 2015, including and especially, regarding the work here (in alphabetical order): J. Dickinson, R. Kimball, I.J. Lovette, S.C.R. Smith, L.M. Stenzler, A.K. Townsend, and R. van den Bussche. We are also extremely grateful to Denise M. Woods for Appendix II, Kevin McGowan for Poconos-related thinking, and a couple of anonymous reviewers of past drafts of this work.

This study was carried out in strict accordance with the recommendations in the Guidelines to the use of Wild Birds in Research of The Ornithological Council, and was approved by the Oklahoma State University IACUC (ACUP No: AS50713). Capture and marking of crows occurred under U.S. Department of Interior, U.S. Geological Society, Federal Bird Banding Permit #22165 (Caffrey). Our work was funded in part by grants from The Payne County Audubon Society, and generous assistance from J. and N. Wilhm, and E. and H. Caffrey.

## APPENDIX I. Details

### 1. Individuals known to be breeding for their first time

a. 2001, Group 42 (Table 1): A 3-year old male who usurped his natal territory (NK); his half-sister remained in the group until eggs hatched and two half-brothers remained and contributed to the feeding of nestlings (Appendix A 1a in Caffrey and Peterson 2015).
b. 2001, Group 45 (Table 1): A 3-year old male (HE, Appendix A 2e in Caffrey and Peterson 2015) who budded a territory off from the group into which he had immigrated. He bred with a female of unknown origin and unrelated to local group members.
c. 2002, Group 40 (Table 1): A 4-year old female (TF; Appendix A 2i in Caffrey and Peterson 2015) who budded a territory off from the group with which she had been caught (late in her first year; she was the daughter of the female breeder but *not* the male). She bred with a male of unknown origin that had immigrated into her group, as an adult, the previous year.

### 2. Nest Building Notes

a. Pair 57 (2002) - a new pair, formed late in the season (end of March) after the death of the male’s previous (and long-time) mate (killed while incubating eggs in a *second* nest [Appendix A 2c in Caffrey and Peterson 2015]) – completed their nest in about five days (although the female added to the nest lining the day before she began incubation) and were intermittently present during checks of their nest area and territory for the following three days until incubation began (Fig. 1g).
b. Pair 115, in 2001, worked intermittently on their first-attempt nest over at least 16 days (Fig. 1d), during which they were also sometimes observed seemingly casually foraging in their territory (often with other group members) or sometimes could not be found, sometimes for half-days at a time (more than once we wondered if this nest had been abandoned). (Group 115’s pattern of absences *during* nest building was unusual for our population.) The female began incubation two days after a watch during which she brought to the nest and worked in a stick, and then abandoned the attempt after 12 days in response to human disturbance (municipal tree-trimmers in surrounding and immediate areas for two days). By three days later their second attempt was underway and the following day at 0640 CST, one of us (CC) watched for one hour as the pair made 25 trips to the nest; 23 of them with nesting material (Fig. 1a, b, and d). They ceased working on the nest by two days later, then avoided the nest area and seemingly casually hung around with other group members in regularly-frequented areas of their territory for two days, after which they were sporadically seen near, at, and in the nest for another two days before the female began incubating eggs.
c. Extra-pair individuals that got into nests appeared to enjoy moving around and repositioning themselves, sometimes adjusting nesting material. In one case, an adult male auxiliary (OM; Appendix A 1b in Caffrey and Peterson 2015) was observed shaking and ruffling up his wings while in sitting in a nest. In field notes (CC): “He looks as if he’s getting a buzz from it!” Sometimes auxiliaries got in nests when alone and sometimes when breeders were present. An example of the latter: a yearling female auxiliary landed on the nest rim while her social half-brother (the breeder) was working on it from inside; she quietly vocalized, and then got in as he got out and onto the rim. He stayed for one minute and left as she sat, repositioned herself a few times, and played a bit with some sticks near the rim for two more minutes before leaving.

### 3. Auxiliary nests

It was not uncommon for auxiliaries to begin, work on, and in some cases, complete nests separate from those of breeding pairs. In 2001 and 2002, we knew of five such nests, those of:

a. KP (Appendix A 1b in Caffrey and Peterson 2015).
b. TM in both years (and 2000); Appendix A 2d in Caffrey and Peterson 2015.
c. FX (Appendix A 2b in Caffrey and Peterson 2015).
d. RA and EK: Early in the nesting season of 2002 (mid February), RA (Group 7, Appendix A 2c in Caffrey and Peterson 2015) and EK (Group 42, Appendix A 1a and 2j in Caffrey and Peterson 2015) were seen together a couple of times in a relatively unused (by crows) area in an overlap zone between Groups 42 and 10 (Fig. 1 in Caffrey and Peterson 2015; boundaries slightly different in 2002); shortly thereafter and although both were “pre-hatch” auxiliaries (Caffrey and Peterson 2015) in their respective groups, they also began, intermittently worked on (over about a week), and ultimately abandoned a nest in this area. The incipient nest had appeared “sloppy” to us, and all evidence of it surprisingly disappeared within days of RA and EK not being seen in its area anymore.

### 4. Incubation Notes

a. Auxiliaries that fed females included yearling and older females and males; these individuals fed their social mothers (N = 15), their genetic mothers (N=2 “Uncertain”s; one individual in both years [FX; Appendix A 2b in Caffrey and Peterson 2015]), their older half-brother’s mate (N = 2), and female breeders of groups into which they immigrated (N = 3; all adult males). One adult male auxiliary (NX) fed a replacement for the female breeder in the group (#333) in which his relationship to breeders had been uncertain (and with whom he fathered two of the four fledglings from this nest; Appendix A 1b, and Appendix B Predictions 8 and 12-15, in Caffrey and Peterson 2015), and one yearling was unmarked and so its relationship with the female breeder was unknown.
b. Auxiliaries not observed to feed incubating females included yearling and older females and males; these individuals did not feed their social mothers (N = 21), their genetic mothers (N=3 “Uncertain”s; Caffrey and Peterson 2015), their older half-brother’s mate (a yearling female and an adult male; the latter was denied access to the incubating female [EK in 2002; above, and Appendix A 2j in Caffrey and Peterson 2015]), females that had replaced their social mothers (N = 4; two adult females and two adult males), the female that had replaced his mother (he was denied access to her [PH; above, and Appendix A 2c in Caffrey and Peterson 2015]), the female breeder in the group when one “Uncertain” was caught (a female for whom we lacked DNA), his older brother’s mate, the mates of males that adult male auxiliaries used to live with (n = 2), and female breeders of groups into which they immigrated (N = 5; three adult females and two adult males).

### 5. Copulation Notes

a. Four days before she began incubation, as she landed on a branch together with her mate, the female lowered her head, spread her wings, and quivered her tail in a precopulatory display (Verbeek and Caffrey 2002) before her mate mounted her. Afterward, they sat together for at least 11 minutes, and during the first one, the female repeatedly (every 5-8 sec) slightly raised and lowered her wings.
b. The female got onto the rim at her mate’s arrival at 1549 CST on Day 3 of incubation.
c. The female stayed in the nest at 943 CST on Day 6.
d. The same female (c) flew to a nearby branch to join her mate at 1427 CST on Day 7.
e. At one point during a watch the male (NK; Appendix A 1a in Caffrey and Peterson 2015) arrived at the nest and the female (YL) raised up from inside, and as NK mounted her, the adult male auxiliary (EK; Appendix A 2j in Caffrey and Peterson 2015) flew in and landed on a branch right above them. NK dismounted to chase EK.
f. Later during the same watch (e), NK arrived - to copulate with YL - at the same time as did the one-year old male auxiliary (having just filled his esophageal pouch with food and water, presumably intended for the female; his mother); NK mounted YL and was flapping vigorously to maintain balance when the one-year old hopped on his back. With the one-year old on top and flapping as well, EK arrived and attempted to slide underneath NK, at which point YL broke loose from the crowd. All four of them then briefly sat together beside the nest, until NK displaced EK to a branch nearby and YL got back in.

### 6. Territorial Behavior

Once pairs of crows selected nest trees and began working on nests, and for much of the rest of the year, defense of territories by most groups consisted of seemingly casual acknowledgment of sometimes seemingly sloppy and sometimes distinct boundaries established previously; group members occasionally called from territory peripheries, and chased most extra-group crows that ventured too close to active nesting areas, but also sometimes joined with neighbors to harass predators or intruders (the latter especially during winter months of pecan fruiting).

## APPENDIX II. Characteristics of American Crow nests, Stillwater OK, 2000

| NEST # | TREE SPECIES | TREE HEIGHT (m) | NEST HEIGHT (m) | NEST PLACEMENT IN TREE | NEST EXPOSURE (visibility<sup>1</sup>) | NEST SOLAR EXPOSURE | SURROUNDING HABITAT |
| --- | --- | --- | --- | --- | --- | --- | --- |
| 1a | Pine | 13.50 | 9.50 | Center, against trunk | 4 (a bit all around) | Partial shade | Park with scattered trees, along path |
| 1b | Pine | 7.40 | 6.40 | Center, top | 5 | Full sun | Park with scattered trees, along path |
| 2 | Pine | 14.40 | 12.60 | Center, top | 4 (top) | Full sun | Park, pine forest patch |
| 3b | Pine | --- | --- | --- | 3 (top, a bit all around) | Full sun | Forest strip along lake and lawn, in back yard |
| 4b | Pine (Scotch) | 9.00 | 8.40 | Center, top | 4 (top, 2 sides) | Full sun | Park, scattered trees |
| 6 | Snag | --- | --- | Center | 4 (all sides, not above) | Full shade | Riparian belt along creek, over water |
| 7 | Oak (red) | 11.70 | 10.98 | Center, against trunk | 2 | Full shade | Treed residential area |
| 10 | Magnolia | 9.15 | 6.30 | Center | 4 (top, 2 sides) | Full sun | Magnolias scattered in field, near construction |
| 11 | Pine | 14.40 | 13.60 | 5 ft. out on limb | 4 | Full sun | Residential street |
| 12 | Red cedar | 11.40 | 9.80 | Center | 4 | Full sun | Field of cedars |
| 14 | Pecan | 23.60 | 17.20 | Crotch of one of several trunks | 3 | Full shade | Field and woods, back yard |
| 15 | Pecan | --- | --- | 5 ft. out on limb | 2 | Full shade | Mixed woodland |
| 17 | Pine | --- | --- | --- | 3 | Partial shade | Back yard, near grass/savannah (x-country track) |
| 19 | Cottonwood | 23.40 | 16.35 | 6 ft. out on limb | 2 (top covered) | Partial shade | Riparian belt along creek edge, lawn, back yard |
| 21 | Red cedar | --- | --- | Center | 1 | Full shade | Fence row, backyard |
| 22 | Red cedar | --- | --- | Center | 1 | Full shade | In field, scattered cedar |
| 25 | Pine | 22.80 | 18.90 | Near trunk | 3 | Partial shade | Front yard, near street |
| 27 | Red cedar | 16.25 | 14.25 | Center | 1 | Full shade | Front yard, near street |
| 28 | Pine | 17.85 | 14.00 | 6 ft. out on limb | 5 | Full sun | Back yard, near street |
| 29 | Pine | 17.00 | 12.15 | Center, top of 1 of 2 trunks | 4 | Full sun | Lone tree, parking lot |
| 32 | --- | --- | --- | --- | 1 | Full shade | Dense forest patch |
| 33 | Red cedar (female) | 11.04 | 7.80 | 7 ft out on limb | 4 | Full sun | Lone tree, field edge, near construction |
| 34 | Elm | --- | --- | Against 1 trunk at "Y" | 1 | Full shade | Back yard |

<sup>1</sup> Visibility of nest to observers on the ground. Scale 1 to 5, 1=completely hidden/ not visible from any direction, 3=visible from some directions, 5=visible from all directions.

## Notes

https://www.caroleecaffrey.com

### Competing Interest Statement

The authors have declared no competing interest.

## LITERATURE CITED

Arnold, K. E., S. L. Ramsay, C. Donaldson, and A. Adam. 2007. Parental prey selection affects risk-taking behavior and spatial learning in avian offspring. Proceedings of the Royal Society B 274:2563–2569.

Caffrey, C. 2002a. Catching crows. North American Bird Bander 26:137–145.

Caffrey, C. 2002b. Marking crows. North American Bird Bander 26:146–150.

Caffrey, C. 2002c. Marking nestling crows: an addendum. North American Bird Bander 27:12.

Caffrey, C. and C. C. Peterson. 2015. Group composition and dynamics in American Crows: insights into an unusual cooperative breeder. Friesen Press. www.caroleecaffrey.com.

Caffrey, C., C. C. Peterson, and T. W. Hackler. 2016. Nestling-care decisions by cooperatively-breeding American Crows. ACB. www.caroleecaffrey.com.

Clark, L. 1987. Thermal constraints on foraging in adult European Starlings. Oecologia 71: 233–238.

Emlen, J. T. 1936. Age determination in the American Crow. Condor 38:99–102.

Gill, S.A., and B.J.M. Stutchbury. 2005. Nest building is an indicator of parental quality in the monogamous neotropical Buff-breasted Wren (Thryothorus leucotis). Auk 122:1169–1181.

Helm, B., T. Piersma, and H. Van Der Jeugd. 2006. Sociable schedules; interplay between avian seasonal and social behavior. Animal Behaviour 72:245–262.

Kalinowski, S. T., M. L. Taper, and T. C. Marshall. 2007. Revising how the computer program CERVUS accommodates genotyping error increases success in paternity assignment. Molecular Ecology 16:1099–1106.

Mainwaring, M.C., and I.R. Hartley. 2013. The energetic costs of nest building in birds. Avian Biology Research 6:12–17.

Moreno, J. 2012. Avian nests and nest-building as signals. Avian Biology Research 5:238–251.

Moreno, J., E. Lobato, S. González-Braojos, and R. Ruiz-de Castañeda. 2010. Nest construction costs affect nestling growth: a field experiment in a cavity-nesting passerine. Acta Ornithologica 45:139–145.

Queller, D. C. and K. F. Goodnight. 1989. Estimating relatedness using genetic markers. Evolution 43:258–275.

Soler, J.J., A.P. Møller, and M. Soler. 1998. Nest building, sexual selection and parental investment. Evolutionary Ecology 12:427–441.

Verbeek, N. A. M., and C. Caffrey. 2002. American Crow (Corvus brachyrhynchos). The Birds of North America, Number 647.

Willmer, P. G. 1983. Thermal constraints on activity patterns in nectar-feeding insects. Ecological Entomology 8: 455–469.

